# Intrinsic, dynamic and effective connectivity among large-scale brain networks modulated by oxytocin

**DOI:** 10.1101/2020.04.22.055038

**Authors:** Xi Jiang, Xiaole Ma, Yayuan Geng, Zhiying Zhao, Feng Zhou, Weihua Zhao, Shuxia Yao, Shimin Yang, Zhongbo Zhao, Benjamin Becker, Keith M. Kendrick

## Abstract

The neuropeptide oxytocin is a key modulator of social-emotional behavior and its intranasal administration can influence the functional connectivity of brain networks involved in the control of attention, emotion and reward reported in humans. However, no studies have systematically investigated the effects oxytocin on dynamic or directional aspects of functional connectivity. The present study employed a novel computational framework to investigate these latter aspects in 15 oxytocin-sensitive regions using data from randomized placebo-controlled between-subject resting state functional MRI studies incorporating 200 healthy subjects. Results showed that oxytocin extensively modulated effective connectivity both between and within emotion, reward, salience and social cognition processing networks and their interactions with the default mode network, but had no effect on the frequency of dynamic changes. Top-down control over emotional processing regions such as the amygdala was particularly affected. Oxytocin effects were also sex-dependent, being more extensive in males. Overall, these findings suggest that modulatory effects of oxytocin on both within- and between-network interactions may underlie its functional influence on social-emotional behaviors, although in a sex-dependent manner. Furthermore, they demonstrate a useful approach to determining pharmacological influences on resting state effective connectivity and support oxytocin’s potential therapeutic use in psychiatric disorders.

## Introduction

The evolutionarily highly conserved hypothalamic neuropeptide oxytocin (OXT) is a key modulator of social and emotional behaviors across species (Bakermans-Kranenburg and Van Ijzendoorn, 2013; Bethlehem *et al*., 2013; Wigton *et al*., 2015; Kendrick *et al*., 2017) and there is a wide distribution of regions expressing oxytocin receptors in the human brain (Bethlehem *et al*., 2013; Quintana *et al*., 2019). In line with OXT’s complex behavioral effects, brain imaging studies have demonstrated its intranasal administration can modulate widely distributed neural systems involved in social cognition, including attention, reward, emotion and cognitive control processing (Bakermans-Kranenburg and Van Ijzendoorn, 2013; Bethlehem *et al*., 2013; Wigton *et al*., 2015; Paloyelis *et al*., 2016; Johnson and Young, 2017; Kendrick *et al*., 2017; Mitre *et al*., 2017).

Combining intranasal OXT administration with resting state fMRI (rsfMRI) has gained increasing interest in order to investigate its modulatory effects on intrinsic functional brain organization while controlling for the influence of external stimuli or contexts (review in Seeley *et al*., 2018). Most previous studies focused on intrinsic amygdala networks and consistently reported an increase in amygdala-prefrontal cortex functional connectivity (Sripada *et al*., 2013; Eckstein *et al*., 2017; Kou *et al*., 2019). Additionally, one study has reported differential modulatory effects of OXT on dorsal and ventral striatal functional connectivity with frontal and cerebellar regions engaged in reward, cognitive control and emotional learning (Zhao *et al*., 2018) and another effects on the intrinsic interaction of default mode and attention-related networks (Xin *et al*., 2018). Together these studies suggest that OXT modulates the intrinsic functional interplay among brain regions and networks involved in emotional control, reward processing and attention. However, there are still several unanswered questions that are of great importance to interpreting which neural circuits OXT modulates and how it influences their functions and the interplay between them. Firstly, many studies have employed hypothesis-driven seed-based analyses which focus on specific brain regions/networks of interest and may therefore limit a more systematic and comprehensive understanding of the modulatory role of OXT. Secondly, current resting state studies have assessed functional connectivity by simply averaging scan-length fMRI BOLD signals and have not addressed possible dynamic temporal variations in functional connectivity. A number of studies have demonstrated dynamic temporal variations in functional connections both within and between different networks (Gilbert and Sigman, 2007; Chang and Glover, 2010; Garrett *et al*., 2010; Protzner *et al*., 2010; Sakoğlu *et al*., 2010; Smith *et al*., 2011; Bassett *et al*., 2011, 2013, 2015; Hutchison *et al*., 2013*a, b*; Mueller *et al*., 2013; Calhoun *et al*., 2014; Li *et al*., 2014; Ou *et al*., 2014; Zhang *et al*., 2016; Vidaurre *et al*., 2017; Jiang *et al*., 2018; Yuan *et al*., 2018). While these latter studies have adopted a number of different methodologies, they have consistently revealed the dynamic nature of both spontaneous neural activity (Garrett *et al*., 2010; Protzner *et al*., 2010) and connectivity/interactions among brain regions/networks. These dynamic changes can be characterized as different ‘states’ occurring within the full scan time series (Chang and Glover, 2010; Hutchison *et al*., 2013*b*; Mueller *et al*., 2013; Calhoun *et al*., 2014; Li *et al*., 2014; Ou *et al*., 2014; Zhang *et al*., 2016; Yuan *et al*., 2018) reflecting both flexibility and adaptability in neural systems (Bassett *et al*., 2011, 2013, 2015; Zhang *et al*., 2016). It is possible that altered functional connectivity between brain regions following OXT administration additionally involves an influence on such temporal dynamics. Thirdly, the majority of both task- and resting-state studies reporting the effects of OXT on functional connectivity use Pearson correlation techniques and as such do not infer causal interactions, i.e., ‘effective connectivity’ (Friston, 2011; Smith *et al*., 2012; Mumford and Ramsey, 2014). Furthermore, the Pearson correlation techniques cannot account for the contribution of other regions to an observed functional connectivity change between specific pairs of regions (Zhang *et al*., 2015). In terms of functional effects these limitations are of particular importance for establishing the extent to which OXT is influencing top-down as opposed to bottom-up processing of social stimuli and the precise circuitry involved as well as its directionality.

In order to address these issues and facilitate a systematic and more causal determination of the modulatory influence of OXT on intrinsic brain organization the present study adopted a novel computational functional connectivity framework to assess the intrinsic, dynamic and effective connectivity among large-scale brain networks previously reported to be modulated by intranasal OXT. To faciliate a robust estimation of the effects we pooled resting state fMRI (rsfMRI) data from previously published randomized, double-blind, between-subject, and placebo (PLC)-controlled pharmacological studies investigating effects of intranasal OXT (Geng *et al*., 2018; Ma *et al*., 2018; Zhao *et al*., 2018), resulting in a large sample of healthy male and female subjects (N=200). Based on previous studies (Bakermans-Kranenburg and Van Ijzendoorn, 2013; Bethlehem *et al*., 2013; Wigton *et al*., 2015; Kendrick *et al*., 2017), we defined a total of 15 different regions of interest (ROIs) implicated in key functional domains influenced by OXT (medial (mAmyg) and lateral (lAmyg) amygdala, dorsal anterior (dACC) and posterior (PCC) cingulate, anterior (aINS) and posterior (pINS) insula, medial (mOFC) and lateral (lOFC) orbitofrontal cortex, dorsomedial (dmPFC) and ventromedial (vmPFC) prefrontal cortex, dorsal (DS) and ventral (VS) striatum and ventral tegmental area (VTA)), resulting in a total of 29 nodes by examining all regions (except VTA) independently in both brain hemispheres.

In line with our previous studies (Chen *et al*., 2013; Lian *et al*., 2014; Ou *et al*., 2014), we first employed a computational framework on rsfMRI BOLD signals of the 29 ROIs for each subject to identify discrete temporal states within the time courses of data acquisition. We then assessed the effective connectivity pattern among the 29 regions for each state and clustered representative effective connectivity patterns across all subjects and states within the OXT and PLC treated groups. Based on recent findings suggesting that OXT modulates the temporal dynamics of intrinsic EEG networks (Schiller *et al*., 2019) we evaluated whether OXT influenced the frequency of brain temporal state switching by comparing the number of identified states between different treatment groups. In order to determine the direction of the effects of OXT on the intrinsic communication within the emotion-cognition networks we next compared the treatment groups with respect to clustered representative effective connectivity patterns. Based on increasing evidence for sex-dependent effects of OXT (Rilling *et al*., 2014; Gao *et al*., 2016; Bethlehem *et al*., 2017; Luo *et al*., 2017; Ma *et al*., 2018; Lieberz *et al*., 2019) we finally explored sex-dependent effects of OXT at the level of intrinsic dynamic effective connectivity.

## Materials and Methods

### Participants, image acquisition and data preprocessing

Resting state fMRI data was from 200 healthy, right-handed Han Chinese college students without current or past psychiatric, neurological, or other medical disorders and originally recruited as part of several previous published pharmacological-fMRI studies (Geng *et al*., 2018; Ma *et al*., 2018; Zhao *et al*., 2018). All subjects had abstained from consuming alcohol or caffeine in the 24-hours prior to the experiments and had provided written informed consent. The original studies were all approved by the local ethics committee and in accordance with the latest revision of the declaration of Helsinki. Participants completed questionnaires measuring mood (Positive and Negative Affect Schedule, PANAS) and anxiety (State-Trait Anxiety Inventory, STAI) before taking part in the randomized double-blind, between-subject placebo-controlled pharmaco-resting state fMRI experiments. None of the female subjects included in the study were taking oral contraceptives or in their menstrual period but no attempt was made to control which phase of their menstrual cycle they were in. The participants whose data were included in the current analysis had been randomly assigned to the two treatment groups ((OXT, n=104, 54 males) vs placebo (PLC, n=96, 52 males)) and received either a single intranasal dose of OXT (40 international units (IU) of OXT dissolved in 0.9% saline and glycerin, Sichuan Meike Pharmaceutical Co. Ltd, Sichuan, China – 5 puffs per nostril) or PLC (supplied by the same company and including the same ingredients other than OXT). A 40 IU dose was chosen in the original studies based on findings that this dose had similar or even improved behavioral and neural effects compared with the more commonly used 24 IU dose (Zhao *et al*., 2017; Geng *et al*., 2018; Shin *et al*., 2018).

### Resting state fMRI data acquisition

T1-weighted structural and resting state fMRI (rsfMRI) data were acquired on the same 3T GE MR750 MRI system 45min after intranasal treatment. The major parameters of T1-weighted structural imaging were as follows: FOV = 256 × 256 mm, TR = 5.97 ms, TE = 1.97 ms, flip angle = 9°, slice thickness = 1 mm, slice number = 128. The rsfMRI was acquired using a T2*-weighted EPI sequence when the participants were instructed to keep their eyes closed and relaxed while not falling asleep. The major parameters of rsfMRI were: FOV = 220×220 mm, TR = 2 s, TE = 30 ms, 3 mm isotropic, slice number = 39, 255 whole-brain volumes (8.5 minutes). Cushions were used to help stabilize the participant’s head to help with head motion control. Six-parameter head motion (< 3 mm) and mean frame-wise displacement (FD, < 0.5 mm) (Power *et al*., 2012) were also used as inclusion criteria to control motion-related artifacts in line with previous studies (Zhao *et al*., 2018).

The rsfMRI data was preprocessed using FSL FEAT (https://fsl.fmrib.ox.ac.uk/fsl/fslwiki/FEAT) in line with our previous study (Zhao *et al*., 2018). The first 10 volumes were discarded for scanner equilibration. The preprocessing steps included motion correction, brain extraction, slice-timing correction, spatial smoothing (5 mm FWHM kernel), intensity normalization, and band-pass filtering (0.01∼0.1 Hz). ICA-AROMA was adopted to remove motion-related artifacts (Pruim *et al*., 2015). White matter and CSF mean signals were removed via linear regression. The individual structural images were firstly normalized to MNI space using FSL FNIRT (12 degrees of freedom (DOF)), and the preprocessed rsfMRI images were then non-linearly registered to the normalized corresponding individual structural images in MNI space.

### Definition of regions of interest

Figure 1 provides a flow chart of the computational framework used in the analysis. The 15 chosen ROIs detailed in Table 1 were identified using three brain atlases (Tzourio-Mazoyer *et al*., 2002; Edlow *et al*., 2012; Fan *et al*., 2016). With the exception of the VTA the ROIs were examined independently in both brain hemispheres resulting in a total of 29 non-overlapping regions. Specifically, the VTA region was defined based on a brainstem atlas (Edlow *et al*., 2012) and the vmPFC in line with previous studies (Jung *et al*., 2018) and dmPFC using the AAL atlas (Tzourio-Mazoyer *et al*., 2002). The remaining ROIs were defined using the Brainnetome atlas (Fan *et al*., 2016). The spatial distribution of the 29 regions is presented in Figure 1A, and the names, indices, and MNI coordinates of the 29 regions are listed in Table 1. Averaged preprocessed BOLD signals were extracted for each of the 29 regions.

**Table 1.**
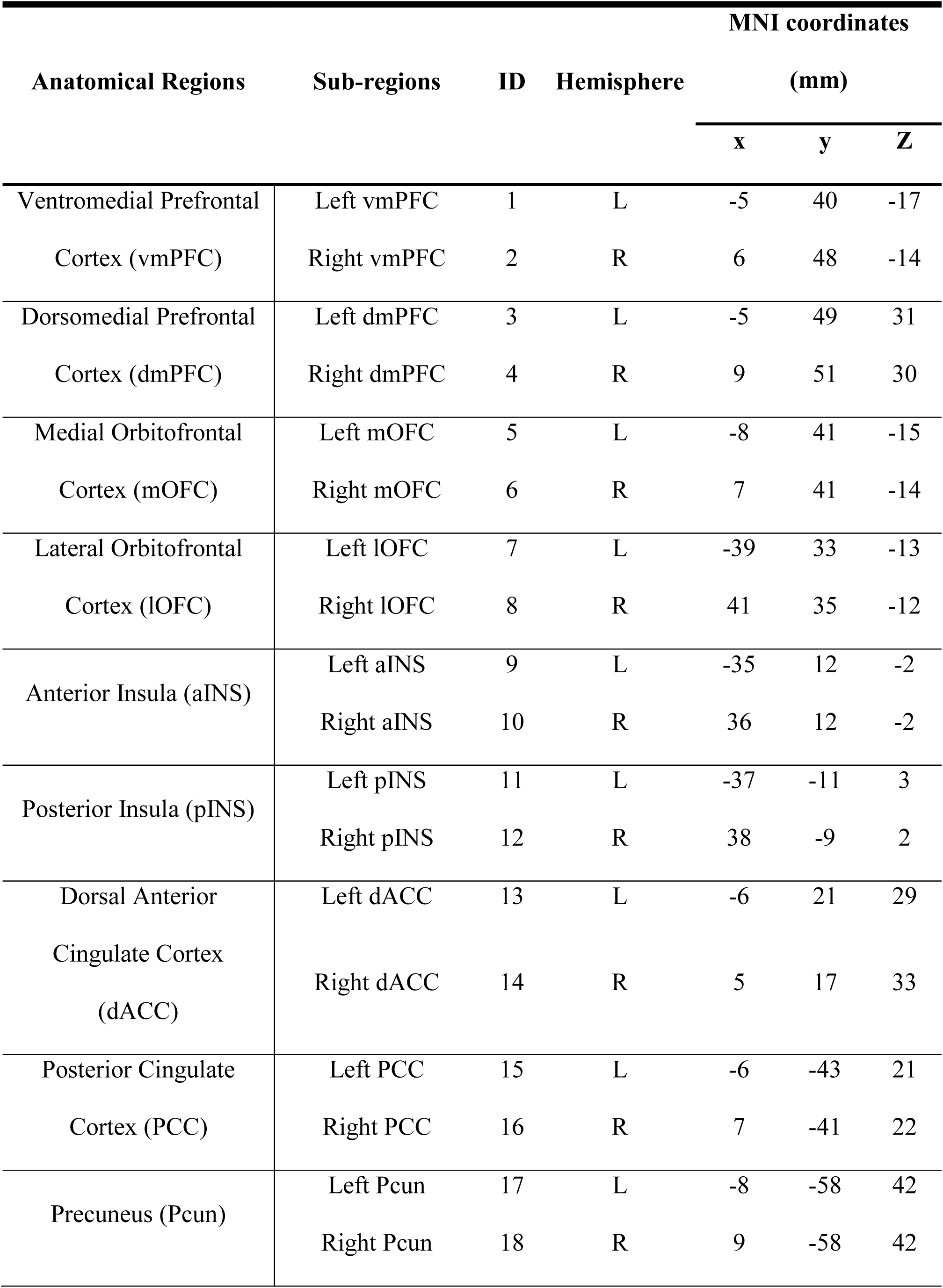
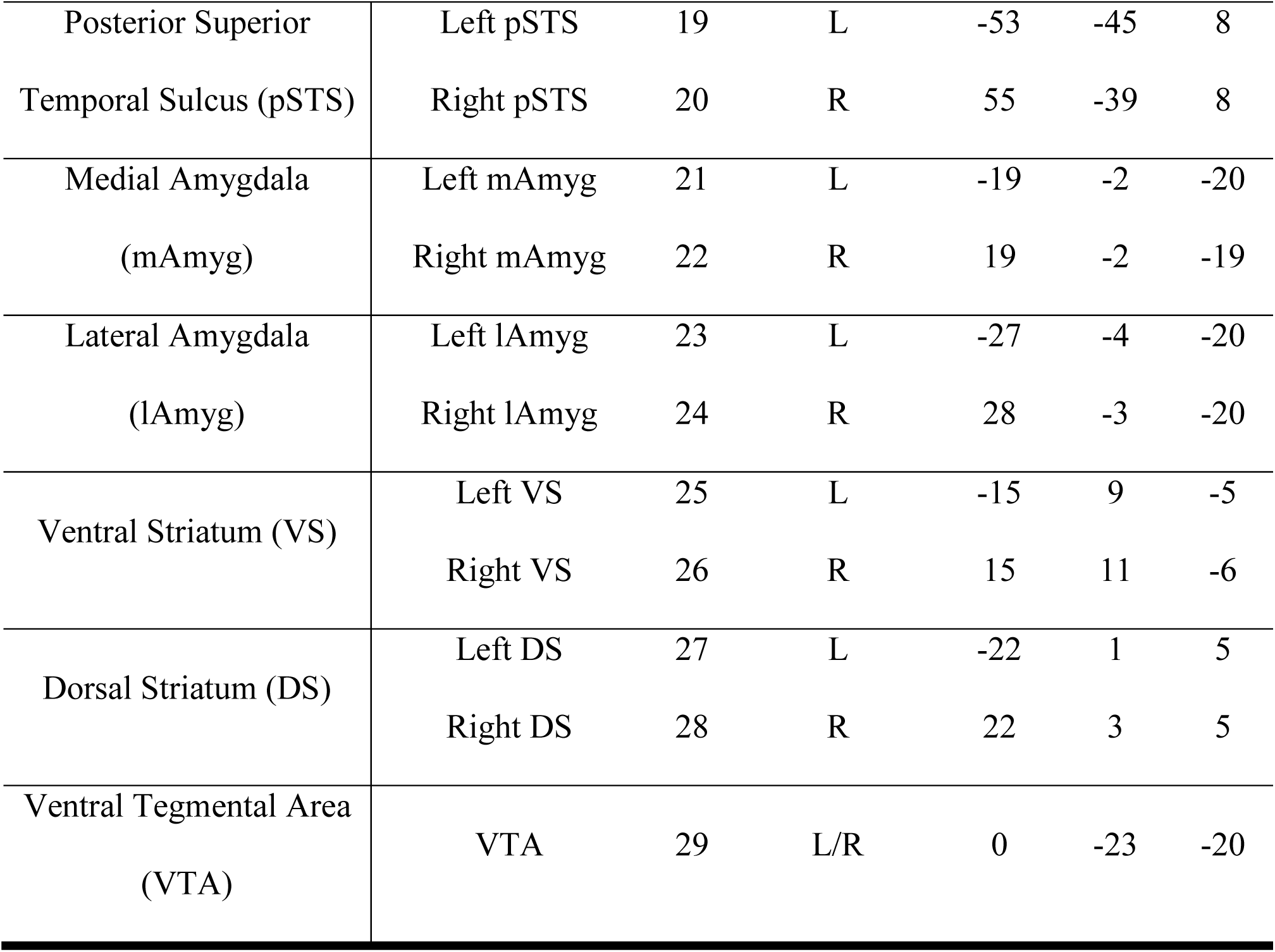
Names, indices, and MNI coordinates of the 29 regions of interest.

**Figure 1.**
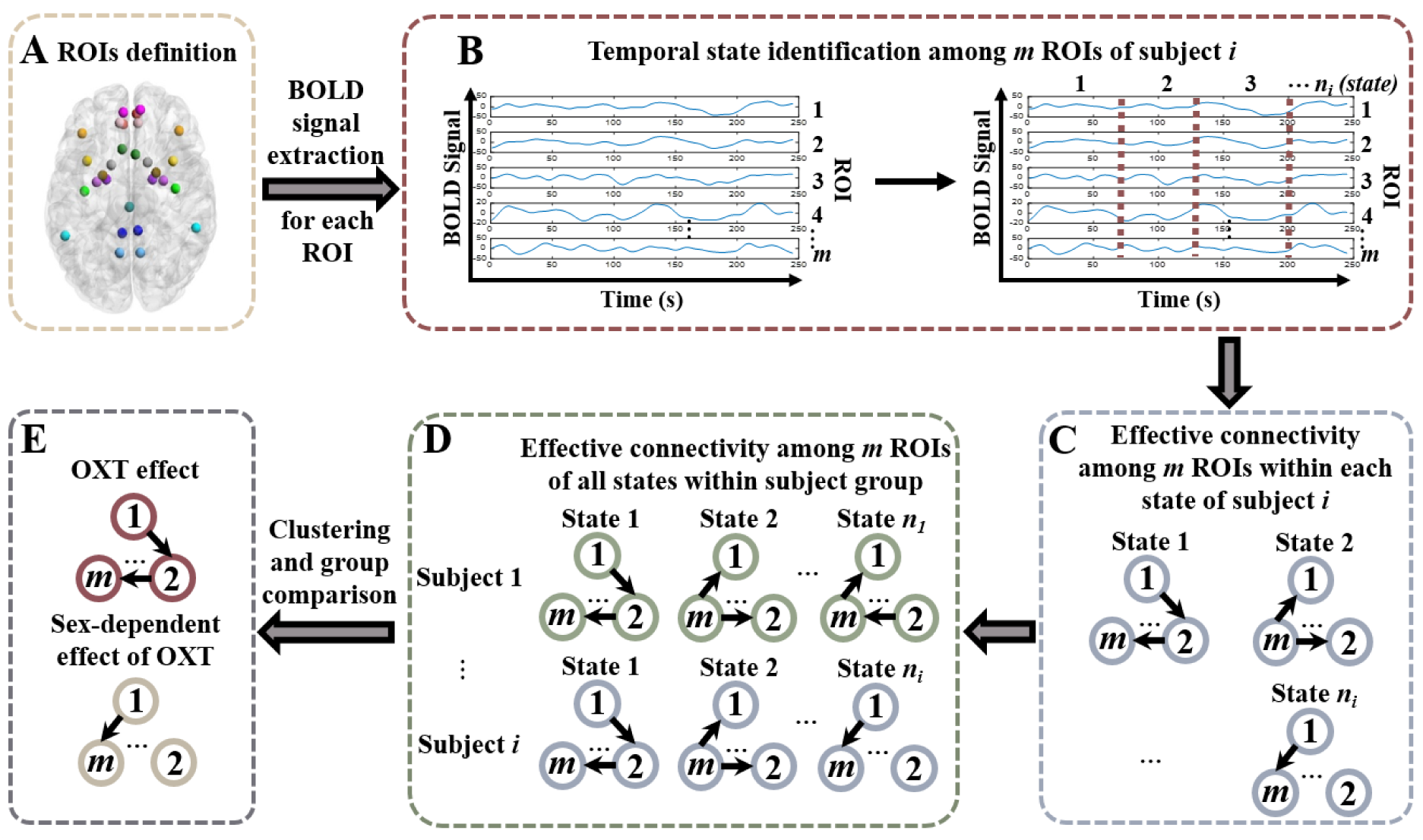
Flow chart of the proposed computational framework. **(A)** Definition of regions of interest (ROIs). **(B)** Temporal state identification of BOLD signals among all *m* ROIs of subject *i*. There are totally *n*_*i*_ states identified for subject *i*. The red dashed lines represent the boundaries of successive states. **(C)** Effective connectivity analysis among all *m* ROIs within each state of each subject *i*. **(D)** Aggregation of effective connectivity among *m* ROIs of all states and subjects within the subject group for clustering and group comparison. **(E)** OXT treatment and sex-dependent effects on temporal dynamic and effective connectivity patterns among *m* ROIs.

### BOLD signal temporal state identification

To characterize temporal dynamics in BOLD signals among the different ROIs (Fig. 1B), we adopted our previous Bayesian connectivity change point model (Lian *et al*., 2014; Ou *et al*., 2014) to identify the change points defined as the abrupt boundaries of temporal states. Conceptually, this model uses a Bayesian inference approach to analyze the joint probabilities of multivariate time series among multiple brain ROIs between different time points and statistically determines the boundaries of the temporal states by means of sampling the posterior probability distribution of each time point as being a change point using a Markov Chain Monte Carlo (MCMC) scheme. The identified change points represented the determined boundaries of temporal states and were indicative of dynamic switching of temporal states among brain regions. To be precise, 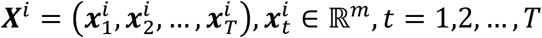 denotes the set of rsfMRI time series of subject *i* (the superscript subject index *i* is omitted in the following equations since we do not refer to any specific subject), *T* is the number of time points, *t* is the time point index, and *m* is the number of brain ROIs. We assumed ***X*** follows an *m*-dimensional Gaussian distribution, i.e., ***x***_*t*_∼*N*(*μ*, Σ), where *μ* is the *m*-dimensional mean vector, and *Σ* is the *m* × *m* covariance matrix. The joint probability of ***x***_1_, ***x***_2_, …, ***x***_*T*_ was calculated as follows:

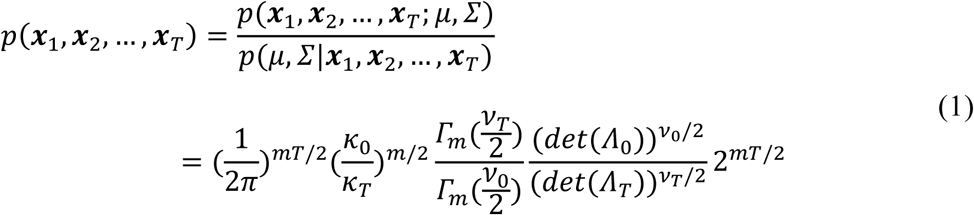

where the conjugate prior distribution and posterior distribution of (*µ*, ∑) is the *N* − *Inv* − *Wishart*(*μ*_0_, *Λ*_0_/*k*_0_, *v*_0_, *Λ*_0_) and the *N* − *Inv* − *Wishart*(*μ*_*T*_, *Λ*_*T*_/*k*_*T*_, *v*_*T*_, *Λ*_*T*_), respectively, and 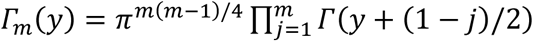 is the multivariate gamma function.

We then defined a change point indicator vector 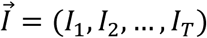 where *I*_*t*_ = 1 if *t*-th time point ***x***_*t*_ was a change point and *I*_*t*_ = 0 otherwise. Therefore there were 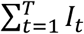 change points in total dividing the whole time series into 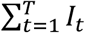 temporal states (assuming *I*_1_ is always the change point and equals 1). We randomly initialized a change point indicator vector 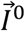 and calculated the posterior distribution 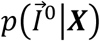 as follows:

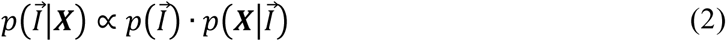

where 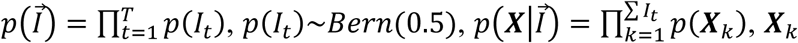 is the rsfMRI time series belonging to the *k*-th temporal state, and *p*(***X***_*k*_) was calculated via Eq. (1). Then for *q*-th iteration (*q=1*,..,*Q*), a new indicator vector 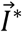 was generated by changing the value of a randomly selected indicator in 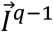 from 0 to 1 or from 1 to 0, and 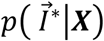 was calculated via Eq. (2). For a randomly generated value *w*∼*uniform*(0,1), if 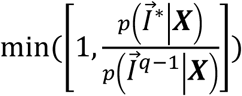, we set 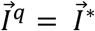, otherwise 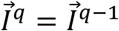. The iteration number *Q* was set large enough (*Q*=5000 in this study) to ensure the convergence of 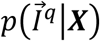. Finally, after removing the burn-in from the MCMC samples of the posterior distribution, we calculated the posterior probability for each time point to be a change point. The number of identified states within the full scan time series may reflect the frequency of brain temporal-varying dynamic switching (Chang and Glover, 2010; Hutchison *et al*., 2013*b*; Mueller *et al*., 2013; Calhoun *et al*., 2014; Li *et al*., 2014; Ou *et al*., 2014; Zhang *et al*., 2016; Yuan *et al*., 2018) and we therefore evaluated whether OXT influenced this by comparing the number of identified states in the different treatment groups.

### Effective connectivity analysis within each temporal state

To investigate the casual relationship among brain ROIs within each of the identified temporal states (Fig. 1C), we performed effective connectivity analysis using the well-established Peter and Clarke (PC) algorithm (Spirtes and Glymour, 1991). Based on the time series of *m* ROIs within a temporal state, the PC algorithm estimated the dependency and directionality among all *m* ROIs (Spirtes and Glymour, 1991). The idea of the PC algorithm is that based on the initial setting of all possible between-region connections, it removes those connections if two regions are (conditionally) independently conditioned on all possible combinations of nodes (multiple comparison corrected) (Spirtes and Glymour, 1991). Next, the direction of each remaining connection was estimated by considering the set of regions causing the conditional independence and the neighbors shared in common by two given regions (Meek, 1995; Mumford and Ramsey, 2014). In line with previous studies (Aliferis *et al*., 2003), we focused on binary information of uni-direction between two ROIs, e.g., ‘1->2’ represents that ROI 1 has an effect on ROI 2 (Fig. 1C).

### Clustering and group-comparison of effective connectivity

To identify overall and sex-dependent effects of OXT on temporal variations in dynamic and effective connectivity among *m* ROIs (Fig. 1E), we pooled the effective connecitvity patterns of all states and subjects within a treatment group (Fig. 1D) and performed clustering in order to obtain clustered representative effective connectivity patterns. The detailed clustering procedure is presented in Figure 2. Each identified effective connectivity pattern among all *m* ROIs within a temporal state of a single subject (Fig. 2A) was represented as an *m*×*m* binary matrix (Fig. 2B). We denoted that the element in the *u*-th row and *v*-th column of the matrix equals 1 if ROI *u* has a causal effect on ROI *v* or 0 if there is no causal effect between ROI *u* and ROI *v*. We vectorized each *m*×*m* binary matrix along columns to an *m*\**m*-dimensional vector (Fig. 2C), and aggregated the vectors of all states and subjects within the treatment group (Fig. 2D). We performed clustering on all aggregated vectors as previously described (Chen *et al*., 2013) to obtain *s* clusters of *m*\**m*-dimensional vectors (Fig. 2E). Note that the optimal number of clusters was also determined in line with our previous work (Chen *et al*., 2013). Each cluster back-transformed as the representative connectivity pattern among all *m* ROIs within the treatment group (Fig. 2F). The overall and sex-dependent effects of OXT on effective connectivity were then assesssed by comparing the clustered representative effective connectivity patterns between different treatment groups. Thus, an identified modulatory effect of OXT on effective connectivity was a clustered representative effective connectivity pattern which only existed in the OXT and not in the PLC group. The sex-dependent effects of OXT on effective connectivity were where OXT differentially modulated clustered representative effective connectivity patterns in males and females.

**Figure 2.**
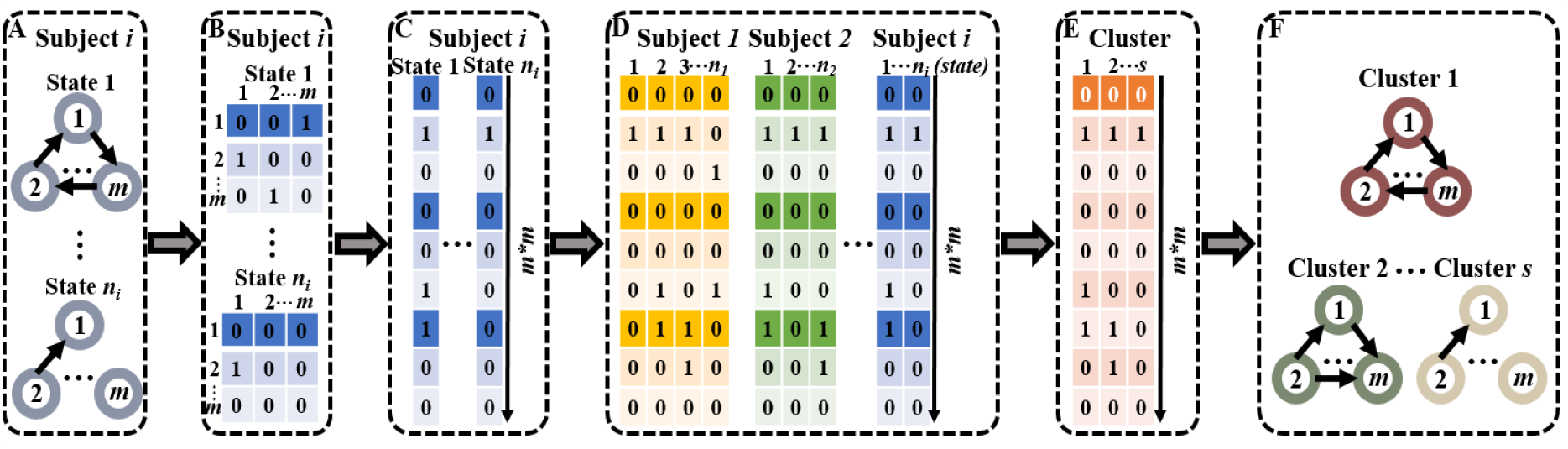
Flow chart of the clustering procedure of time-varying dynamic effective connectivity patterns. **(A)** Identified effective connectivity pattern among all *m* ROIs within each temporal state of a single subject *i*. **(B)** An *m*×*m* binary matrix representing an effective connectivity pattern among *m* ROIs in **(A)**. The element in the *u*-th row and *v*-th column of the matrix equals 1 if ROI *u* has a causal effect on ROI *v*, otherwise 0 if there is no causal effect between ROI *u* and ROI *v*. **(C)** Vectorized *m*\**m*-dimensional vector based on the *m*×*m* binary matrix along columns. **(D)** Aggregated vectors of all states and subjects within a treatment group for clustering. **(E)** Clustered representative vectors. The number of clusters is *s*. **(F)** The representative connectivity patterns among *m* ROIs within a treatment group corresponding to each of the clustered vectors in **(E)**.

## Results

### Potential confounders - head motion, demographics, mood and anxiety

There was no significant difference in mean frame-wise displacement in head motion between the treatment groups (OXT: Mean = 0.10, SD = 0.03; PLC: Mean = 0.10, SD = 0.04; t = -0.33, p = 0.74, independent sample t-test). Furthermore, the treatment groups did not differ in age, gender, anxiety or mood indices as detailed in Table 2.

**Table 2.**
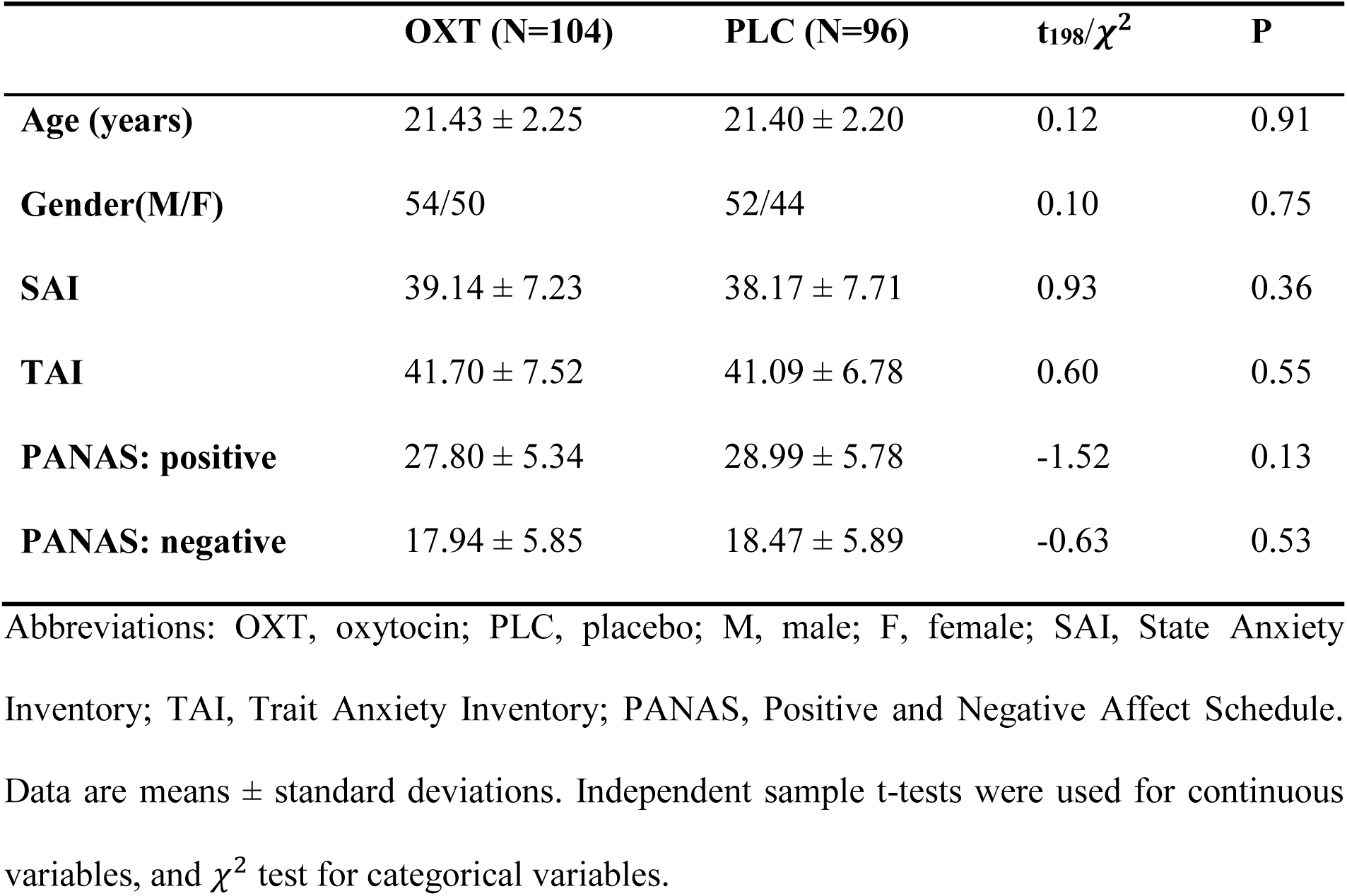
Age, gender, trait and mood indices of the two treatment groups.

|  | **OXT (N=104)** | **PLC (N=96)** | **t<sub>198</sub>/χ<sup>2</sup>** | **P** |
| --- | --- | --- | --- | --- |
| **Age (years)** | 21.43 ± 2.25 | 21.40 ± 2.20 | 0.12 | 0.91 |
| **Gender(M/F)** | 54/50 | 52/44 | 0.10 | 0.75 |
| **SAI** | 39.14 ± 7.23 | 38.17 ± 7.71 | 0.93 | 0.36 |
| **TAI** | 41.70 ± 7.52 | 41.09 ± 6.78 | 0.60 | 0.55 |
| **PANAS: positive** | 27.80 ± 5.34 | 28.99 ± 5.78 | -1.52 | 0.13 |
| **PANAS: negative** | 17.94 ± 5.85 | 18.47 ± 5.89 | -0.63 | 0.53 |

Abbreviations: OXT, oxytocin; PLC, placebo; M, male; F, female; SAI, State Anxiety Inventory; TAI, Trait Anxiety Inventory; PANAS, Positive and Negative Affect Schedule. Data are means ± standard deviations. Independent sample t-tests were used for continuous variables, and χ<sup>2</sup> test for categorical variables.

### Effects of OXT on frequency of brain temporal state switching

We assessed the effects of OXT on frequency of brain temporal state switching by comparing the number of identified temporal states. A two-way ANOVA performed with treatment (OXT, PLC) and sex (male, female) as between subject factors did not reveal either main effects of treatment (d.f. = 1, F = 0.01, P = 0.91) or sex (d.f. = 1, F = 0.19, P = 0.66) or treatment x sex interaction effects (d.f. = 1, F = 1, P = 0.32), suggesting that OXT had no significant effect on frequency of brain temporal state switching.

### Effects of OXT on dynamic effective connectivity

We found a dynamic effective connectivity pattern occurring in the OXT-but not PLC-treated group as shown in Figure 3. The effects of OXT on dynamic effective connectivity of 54 links among the 29 ROIs are detailed in Fig. 3A and summarized in Figure 3B. Overall, the greatest influence of OXT was on causal effective connections from regions in the salience network (23 links from dACC, aINS and pINS) and the posterior midline default mode network (11 links from PCC and Pcun). The predominant effects of OXT on effective connectivity were as follows: 1) Medial and lateral amygdala receive increased effective connections from salience (dACC, pINS), reward (DS, VS and VTA) and social cognition (pSTS) networks, but not from the midline default mode (dmPFC, vmPFC, PCC and Pcun) or directly from aINS, mOFC or lOFC. OXT did not influence any effective connections from the amygdala; 2) increased effective connectivity within the salience network in a rostro-caudal direction (dACC>aINS>pINS; 3) increased effective connectivity from the posterior part of the midline default network (PCC and Pcun) to the salience network (dACC and aINS); 4) increased reciprocal effective connectivity between prefrontal and orbitofrontal cortex regions and aINS, with the same frontal regions also influencing the pINS; 5) increased effective connectivity from pSTS, pINS and PCC to brain reward regions (DS, VS and VTA (only pSTS)); 6) increased homotopic interhemispheric effective connectivity in the majority of regions (12/14 – exceptions are amygdala and pSTS).

**Figure 3.**
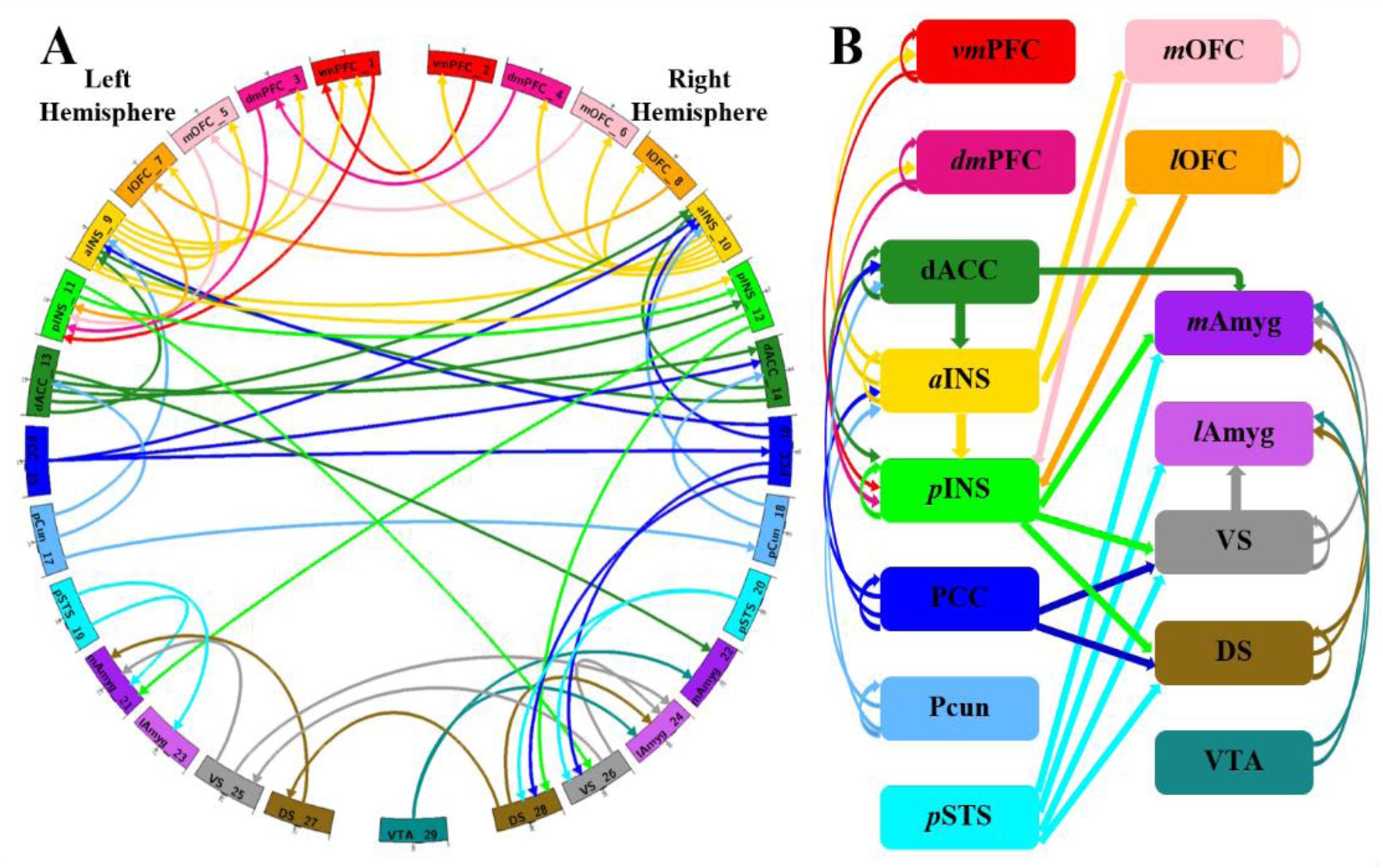
Effects of OXT on dynamic effective connectivity. **(A)** OXT-modulated changes in dynamic effective connectivity among all *m* (*m* = 29 – see **Table 1**) ROIs. **(B)** Overall summary of OXT modulated dynamic effective connectivity patterns among the 15 key brain regions. Arrows denote the direction of causal effects and are color coded by the region of origin. mAmyg = medial amygdala, lAmyg = lateral amygdala, dACC = dorsal anterior cingulate, PCC = posterior cingulate cortex, aINS = anterior insula; pINS = posterior insula, mOFC = medial orbitofrontal cortex, lOFC = lateral orbitofrontal cortex, Pcun= precuneus, dmPFC = dorsomedial prefrontal cortex, vmPFC = ventromedial prefrontal cortex, DS = doral striatum, VS = ventral striatum, VTA = ventral tegmental area.

### Sex-dependent effects of OXT on dynamic effective connectivity

A number (n = 15 links) of OXT-modulated dynamic effective connectivity patterns were found in males but not females (see Figure 4) suggesting that its effects are sex-dependent. There were no OXT-modulated patterns specifically associated with females. The main OXT-modulated patterns found only in males were as follows: 1) increased effective connectivity within the salience network (dACC, aINS and pINS – particularly dACC), and between the posterior midline default network and the dACC; 2) increased effective connectivity between VTA and lAmyg; 3) particularly homotopic interhemispheric connections in orbitofrontal cortex, salience network (dACC, aINS), posterior midline default mode network (PCC, Pcun) and DS. Effective connectivity which was influenced equivalently in the two sexes therefore included: 1) all the effective connections to the amygdala from salience (dACC, pINS), reward (DS, VS and VTA - other than lAmyg) and social cognition (pSTS) networks; 2) all effective connectivity involving the dmPFC and vmPFC and mOFC and lOFC; 3) effective connectivity between aINS and pINS; 4) effective connectivity from the posterior part of the midline default network (PCC and Pcun) to part of the salience network (only aINS); 4) all effective connectivity between pSTS pINS and PCC and striatal reward regions; 5) homotopic interhemispheric connections in dmMPFC, vmPFC, pINS and VS. There were no sex differences between patterns of effective connectivity in the PLC group.

**Figure 4.**
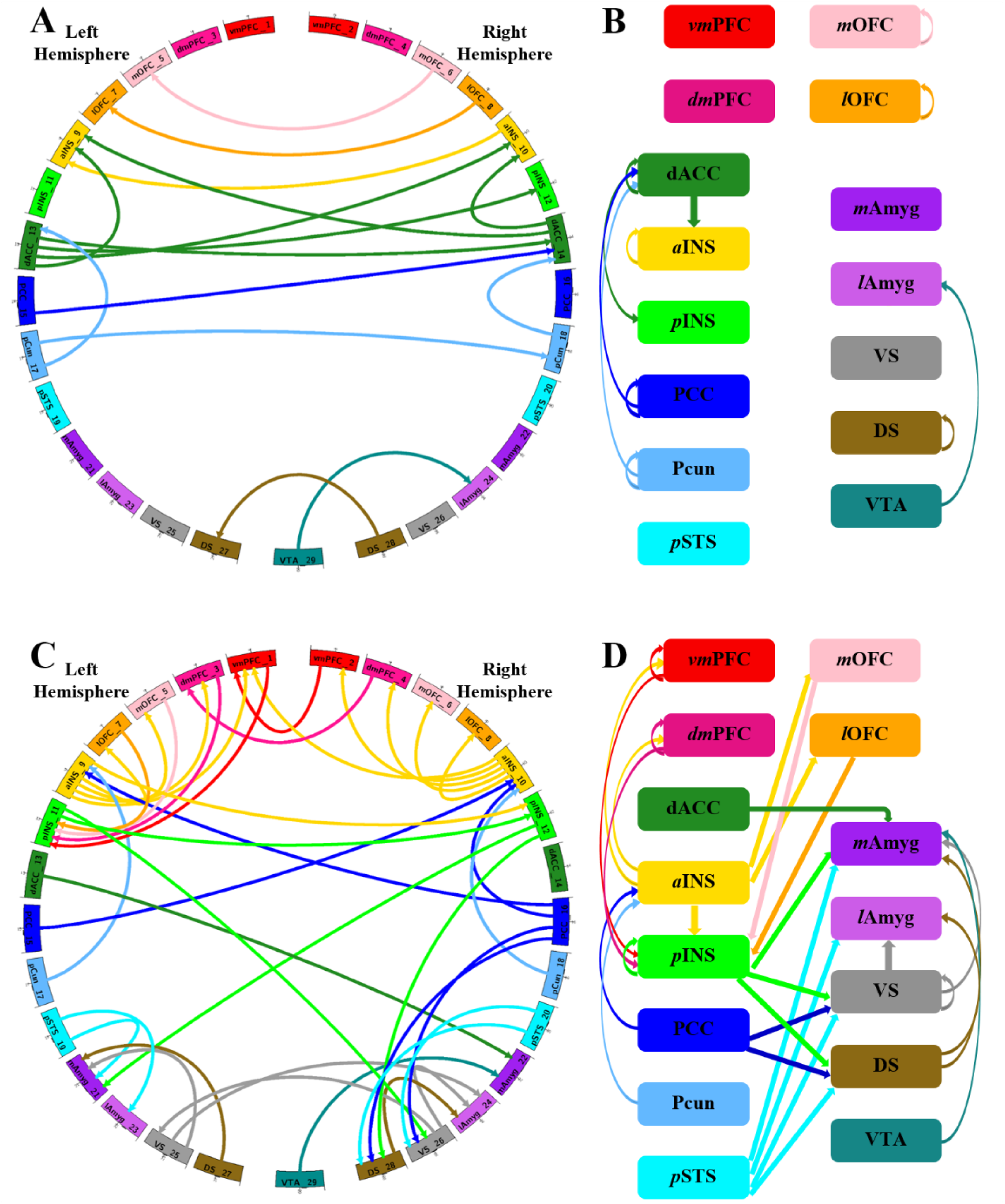
Differential effects of OXT on dynamic effective connectivity patterns in males and females. **(A)** OXT-modulated dynamic effective connectivity changes among all *m* (*m* = 29 – see **Table 1**) ROIs occurring in males but not in females. **(B)** Summary of OXT-modulated dynamic effective connectivity pattern among the brain regions in males but not females. **(C)** Equivalent effective connectivity changes induced by OXT in males and females **(D)** Summary of effective connectivity patterns induced by OXT in males and females. Arrows denote the direction of causal effects and are color coded by the region of origin. mAmyg = medial amygdala, lAmyg = lateral amygdala, dACC = dorsal anterior cingulate, PCC = posterior cingulate cortex, aINS = anterior insula; pINS = posterior insula, mOFC = medial orbitofrontal cortex, lOFC = lateral orbitofrontal cortex, Pcun= precuneus, dmPFC = dorsomedial prefrontal cortex, vmPFC = ventromedial prefrontal cortex, DS = doral striatum, VS = ventral striatum, VTA = ventral tegmental area.

### Reproducibility and classificability of OXT-modulated changes in effective connectivity

We assessed the reproducibility of OXT-modulated effective connectivity across different subjects by counting the number of the OXT-modulated connectivity patterns within each subject. Overall, 80.76% of subjects in OXT group exhibited the corresponding pattern (Fig 3). For the OXT modulated effective connectivity pattern in males but not females (Fig. 4A-B), 83.33% of male subjects exhibited such a pattern. Together, these results suggest reasonable reproducibility of OXT-modulated effective connectivity across subjects. Furthermore, a support vector machine (SVM) analysis demonstrated that the OXT-modulated effective connectivity pattern achieved a promising classification accuracy of 79.81% for distinguishing OXT from PLC group members, and sex-dependent effects achieved an 81.48% accuracy for distinguishing between OXT treated males and females.

## Discussion

The current study employed a novel computational functional connectivity framework to investigate the modulatory effects of the neuropeptide OXT on temporal state switching frequency and the directionality of resting state functional connectivity in brain regions previously shown to exhibit sensitivity to its intranasal administration. While results did not reveal an influence of OXT on temporal state switching frequency they did demonstate widespread (n = 54) changes in effective connectivity involving the 15 ROIs which were primarily characterized by increased connectivity both within and between different functional networks, and notably only top-down influences on the amygdala. Additionally, OXT-evoked changes showed a marked sex-dependency, being more extensive in males than in females.

### OXT effects on brain temporal state switching frequency

Brain temporal state switching frequency may reflect brain flexibility (Bassett *et al*., 2011, 2013, 2015, Zhang *et al*., 2016), i.e., the higher the frequency of brain temporal state switching is, the more flexible the brain is. In support of this conceptualization higher temporal flexibility of the brain has been associated with the motor-skill learning ability (Bassett *et al*., 2011, 2013, 2015) and general intelligence (Zhang *et al*., 2016) and has been found to be impaired across psychiatric disorders characterized by cognitive and social impairments (Zhang *et al*., 2016). However, in contrast to a previous pharmacological EEG study (Schiller *et al*., 2019), we found no evidence of OXT influencing brain temporal state switching frequency. This may perhaps reflect the fact that the peptide has no reported effects on either motor learning or general intelligence, although since it does influence learning with social feedback and rewards (Hurlemann *et al*., 2010; Hu *et al*., 2015) and it is possible that it might influence switching frequency during performance of such social learning tasks. Moreover, the higher temporal resolution of EGG might be more sensitive to determine subtle effects on dynamic switching of brain states.

### OXT effects on dynamic effective connectivity among networks

In line with our hypotheses we observed extensive connectivity changes following OXT administration involving all of the 15 chosen ROIs indicating that it exerts a broad regulatory influence on intrinsic brain functional communication both within and between in attention, salience, reward, social cognition and default networks. While a number of previous resting state studies using conventional Pearson correlation or independent component analysis approaches have tended to emphasize effects of OXT on altered functional connectivity between rather than within brain networks (Xin *et al*., 2018) our current findings indicate that it also has a strong influence on processing within networks.

One of the current main hypotheses concerning the fundamental modulatory actions of OXT is that it increases the salience of social stimuli in a context dependent manner (see Shamay-Tsoory and Abu-Akel, 2016). In support of this hypothesis, a notable finding in our current study is that OXT predominantly strengthens effective connections within the salience network (dACC, aINS and pINS - Menon and Uddin, 2010). The dACC and insula are of particular importance in cognitive evaluation and reappraisal (Picó-Pérez *et al*., 2019) and many reported effects of OXT involve its influence on social decision making and judgments (Gao *et al*., 2016; Kendrick *et al*., 2017; Xu *et al*., 2019). We also found that OXT increased the causal influence of the anterior on the posterior insula supporting findings from our previous task-based study demonstrating that OXT may facilitate switching between interoceptive and external salience processing within the insula (Yao *et al*., 2018a). Furthermore, OXT strengthened reciprocal interactions between the salience network and medial frontal and orbitofrontal cortex regions involved in emotional and cognitive control. This confirms increasing evidence for an influence of OXT on both bottom up and top down control of social salience processing (Yao *et al*., 2018a, 2018b; Xu *et al*., 2019). Additionally, OXT strengthens effective connections from the pINS to basal ganglia reward regions (DS and VS) and those involved in emotion processing (amygdala) suggesting a potential mechanism via which OXT can facilitate reward and emotional responses to socially salient stimuli. Decreased functional connectivity within the salience network is a hallmark of both clinical (Ebisch *et al*., 2011) and trait (Di Martino *et al*., 2016) autism and the present findings further support the proposed therapeutic potential of OXT to normalize neural and behavioral dysregulations in this disorder (see Kendrick *et al*., 2017).

In agreement with a recent independent component analysis of OXT-evoked resting state changes using part of the same dataset as in our current study (Xin *et al*., 2018) we observed that OXT strengthened effective connectivity between parts of the default mode and salience networks. Specifically, the main anterior hubs of the midline default mode network (dmPFC and vmPFC) showed increased causal connections with the pINS following OXT whereas the posterior default mode hub (PCC and Pcun) showed increased connections with the aINS and dACC. Interactions between default and salience networks are important for switching attention between internal and external cues (Seeley *et al*., 2007; Uddin *et al*., 2009) and are strongly implicated in social-cognitive and self-referential processes (Schilbach *et al*., 2008; Mars *et al*., 2012). Previous intranasal OXT intranasal administration studies have reported effects in these domains (Zhao *et al*., 2016, 2017; Kendrick *et al*., 2017).

A considerable number of studies have implicated the amygdala as one of the key sites of functional effects of intranasal OXT (Gamer *et al*., 2010; Gao *et al*., 2016; Spengler *et al*., 2017; Kou *et al*., 2019; Lieberz *et al*., 2019), particularly in the context of the processing and learning of emotional cues. These studies have been unable to determine whether OXT primarily influences bottom up (i.e. effective connections from) or top-down (effective connections to) amygdala control since causality could not be established. This is of particular importance since strengthening top-down control over amygdala responses to negative emotional stimuli is considered an important therapeutic strategy in the context of anxiety and depression disorders (Zhao *et al*., 2019). Indeed, several previous resting state studies have reported that intranasal OXT increases the strength of connectivity between the amygdala and frontal cortex in patients with anxiety disorders (Dodhia *et al*., 2014; Koch *et al*., 2016). Our current findings therefore of great importance in this respect by demonstrating that OXT selectively influences top down effective connections targeting the amygdala with contributions from salience (dACC and pINS), reward (DS, VS and VTA) and social cognition (pSTS) networks. There was a notable absence of OXT influencing direct effective connections from medial frontal and orbitofrontal regions on the amygdala despite a number of correlational resting state studies having reported functional connectivity changes in these pathways following OXT (Sripada *et al*., 2013; Fan *et al*., 2014; Ebner *et al*., 2016; Eckstein *et al*., 2017; Kou *et al*., 2019). This may reflect difficulties in assessing direct functional connectivity changes between time-series for specific pairs of brain regions where there can be considerable influences from third party regions when Pearson correlation techniques are used (Zhang *et al*., 2015). Indeed, from the pattern of OXT-evoked effective connectivity changes we have observed it may influence the amygdala indirectly via the pINS in the salience network or even more indirectly via the influence of the latter on striatal reward regions (DS and VS). In support of this, several studies have reported that OXT can increase functional connectivity between the insula and amygdala during social processing (Hu *et al*., 2015; Gao *et al*., 2016) as well as during the resting state (Frijling *et al*., 2016).

A large number of animal model and human studies have demonstrated effects of OXT on brain reward systems in line with its proposed role in enhancing reward processing in social contexts (Kendrick *et al*., 2017). The present study found that OXT increased the causal effects of the VTA and DS and VS reward regions on both the medial and lateral amygdala. In animal models the effects of OXT on the VTA are crucial for reward-like properties of social interactions (Song *et al*., 2016) and OXT in the VTA promotes prosocial behaviors in mice during social interactions by increasing activity in reward-specific dopaminergic VTA neurons which project to the striatum (Hung *et al*., 2017). Human studies investigating effects of intranasal OXT have reported increased activity in brain reward regions including VTA and nucleus accumbens in both males (Scheele *et al*., 2013) and females (Scheele *et al*., 2016) associated with its enhancement of the perceived attractiveness of romantic partners. Increased activation in VTA in response to cues of happy (social reward) and angry (social punishment) faces has also been reported following OXT (Groppe *et al*., 2013) together with increased activity in striatal regions and their functional connectivity with the amygdala in a reinforcement associated learning task with social but not non-social feedback (Hu *et al*., 2015). Intranasal OXT also evokes extensive changes in regional cerebral blood flow in brain reward regions (Paloyelis *et al*., 2016) as well as their resting state functional connectivity (Zhao *et al*., 2018). Importantly in further support of the proposed role of OXT in increasing the reward of salient social stimuli the VTA, DS and VS received increasing causal influence from salience (pINS), social cognition (pSTS) and posterior default mode (PCC) networks following OXT.

Interestingly, one of the strongest effects of OXT on effective connectivity within networks was on homotopic interhemispheric connections with 11 out of 14 regions showing this effect. Interhemispheric integration is important for a number of cognitive and social functions and weakened homotopic functional connectivity has been observed in both autism (Li *et al*., 2019) and schizophrenia (Guo *et al*., 2014). Our findings therefore suggest that OXT may play an important functional role in influencing within network interhemispheric communication.

### Sex-specific effects of OXT on dynamic effective connectivity among networks

We found that OXT modulated a number of dynamic effective connectivity patterns in males but not females suggesting that it has sex-dependent effects. Most notably, in men OXT particularly influenced effective connectivity both within the salience network (dACC to aINS and pINS) and its causal connections from the posterior midline default mode network (PCC and Pcun to dACC) as well as homotopic interhemispheric connectivity in these regions. Additionally, the increased VTA causal influence on the lateral amygdala following OXT was also specific to males. The main implication of this appears to be that in males OXT exerts stronger influence on the dACC than in females suggesting that it may have a greater potential for increasing top-down emotion regulation in males. The dACC and insula are both robustly involved in cognitive evaluation and reappraisal (Picó-Pérez *et al*., 2019) and since OXT can differentially influence social evaluations and decision making in both sexes (Scheele *et al*., 2014; Gao *et al*., 2016; Xu *et al*., 2019) our current findings suggest a potential regulatory mechanism via which OXT may gate sex-differential effects even in the absence of external social stimuli. While a previous resting state study did not observe sex-dependent effects of OXT on functional connectivity between attention, salience and default mode networks (Xin *et al*., 2018), a number of task-related studies have demonstrated opposite effects of OXT on dACC, insula and amygdala responses during perception or judgments of socio-emotional stimuli in males and females (Rilling *et al*., 2014; Gao *et al*., 2016; Luo *et al*., 2017; Ma *et al*., 2018; Lieberz *et al*., 2019). Thus, it is possible that the sex differences in effective connectivity we have found in response to OXT could reflect corresponding behavioral sex differences in processing of social stimuli and associated decision making.

### Methodological considerations

The present study adopted our previous Bayesian connectivity change point model (Lian *et al*., 2014; Ou *et al*., 2014) to identify the temporal states of BOLD signals among brain regions. Existing studies have proposed a variety of methodologies from different perspectives to characterize the temporal-varying dynamics of neural activity as well as functional connectivity/interaction (Gilbert and Sigman, 2007; Chang and Glover, 2010; Garrett *et al*., 2010; Sakoğlu *et al*., 2010; Smith *et al*., 2011; Bassett *et al*., 2011, 2013, 2015; Hutchison *et al*., 2013*a, b*; Mueller *et al*., 2013; Calhoun *et al*., 2014; Li *et al*., 2014; Ou *et al*., 2014; Zhang *et al*., 2016; Vidaurre *et al*., 2017; Jiang *et al*., 2018; Yuan *et al*., 2018). Given the lack of ground-truth for the temporal dynamics of functional connectivity in fMRI, validations on simulated data and cross-subject reproducibility together with seeking associations with individual behavioral information were widely considered as reasonable justifications for the efficacy of those methodologies. Our model has been validated by extensive experiments on both simulated and real fMRI data as detailed in Lian *et al*., 2014 and Ou *et al*., 2014 in order to identify stable and meaningful temporal states of fMRI data. The present study based on pharmaco-rsfMRI data has further established the efficacy of our method in identifying meaningful dynamic effective connectivity patterns which exhibited reasonable reproducibility across a large sample of subjects.

We adopted the widely-used Peter and Clarke (PC) algorithm (Spirtes and Glymour, 1991) to perform effective connectivity analysis. Previous studies have reported consistent fMRI effective connectivity results using PC and other methods such as GES (review in Henry and Gates, 2017). Note that both the PC algorithm and commonly used Granger causality analysis (GCA) (Granger, 1969) were both designed to infer effective connectivity patterns. However, in contrast the GCA, the PC algorithm infers all brain ROIs simultaneously and does not rely on the temporal lag information of the fMRI time series. Thus, the PC algorithm tends to be more robust and reliable for functional effective connectivity estimation given the potential influence of relatively low temporal resolution of fMRI data as well as the intervention of hemodynamics on it (Smith *et al*., 2012; Sun *et al*., 2012; Henry and Gates, 2017).

### Limitations

There are several limitations which should be acknowledged in the current study. A large scaled ROI based analysis was used rather than a whole brain one because the computational power that would be required for a whole brain approach using our analysis methods would have been too great. We also did not take into account a possible contribution of menstrual cycle phase on sex differences, although previous studies investigating effects of intranasal OXT resting state or task-dependent changes in females have not reported such effects (Hu *et al*., 2015; Gao *et al*., 2016; Bethlehem *et al*., 2017; Geng *et al*., 2018; Xin et al., 2018; Lieberz *et al*., 2019; Xu *et al*., 2019).

## Conclusion

To the best of our knowledge the current study is the first to systematically assess the effects of OXT adminsitration on intrinsic, dynamic and effective connectivity among large-scale networks in a large sample of both males and females. While the present findings reveal no significant effects of OXT on brain temporal state switching frequency, they do demonstrate extensive connectivity changes both within and between functional networks, including attentional, emotional, reward, salience and cognitive control processing as well as the default mode network. Furthermore, there are sex-dependent effects of OXT particularly involving the salience network which may underlie the increasing number of task-dependent studies reporting opposite effects of OXT on neural and behavioral responses to socio-emotional stimuli. Our findings should also help inform the development of OXT as a potential therapeutic intervention in neuropsychiatric disorders such as autism spectrum disorder and schizophrenia which are associated with both social-emotional deficits and disrupted interactions among networks (e.g., Modahl *et al*., 1998; Hollander *et al*., 2007; Andari *et al*., 2010; Feifel *et al*., 2010; Guastella *et al*., 2010; Pedersen *et al*., 2011; Bakermans-Kranenburg and Van Ijzendoorn, 2013; Kendrick *et al*., 2017).

